# A Facile Method for Determining Lanthipeptide Stereochemistry

**DOI:** 10.1101/2023.10.26.564210

**Authors:** Youran Luo, Shuyun Xu, Wilfred A. van der Donk

## Abstract

Lanthipeptides are a large group of natural products that belong to the ribosomally synthesized and post-translationally modified peptides (RiPPs). Lanthipeptides contain lanthionine and methyllanthionine bis amino acids that have varying stereochemistry. The stereochemistry of new lanthipeptides is often not determined because current methods require equipment that is not standard in most laboratories. In this study, we developed a facile, efficient, and user-friendly method for detecting lanthipeptide stereochemistry utilizing advanced Marfey’s analysis. Under optimized conditions, 0.05 mg peptide is sufficient to characterize the stereochemistry of five (methyl)lanthionines of different stereochemistry using a simple liquid chromatography set-up, which is a much lower detection limit than current methods. In addition, we describe methods to readily access standards of the three different methyllanthionine stereoisomers and the two different lanthionine stereoisomers that have been reported in known lanthipeptides. The developed workflow uses commonly used non-chiral column system and offers a scalable platform to assist antimicrobial discovery. We illustrate its utility with an example of a lanthipeptide discovered by genome mining.

---

Lanthipeptides comprise one of the largest classes of ribosomally-synthesized and post-translationally modified peptides (RiPPs).^1^ These peptides possess various biological functions, including antibacterial, antiviral, antifungal, and morphogenetic activities.^2-4^ Lanthipeptides are conformationally restrained to recognize their biological targets through thioether crosslinks called lanthionine (Lan) and methyllanthionine (MeLan). These structures are formed by enzymatic dehydration of Ser and Thr residues followed by Michael-type addition of the thiols of Cys residues to the dehydrated amino acids (**Figure 1**).^5^ Lanthipeptides owe their various bioactivities to the intricate stereochemistry embedded in their three-dimensional geometry.

**Fig. 1.**
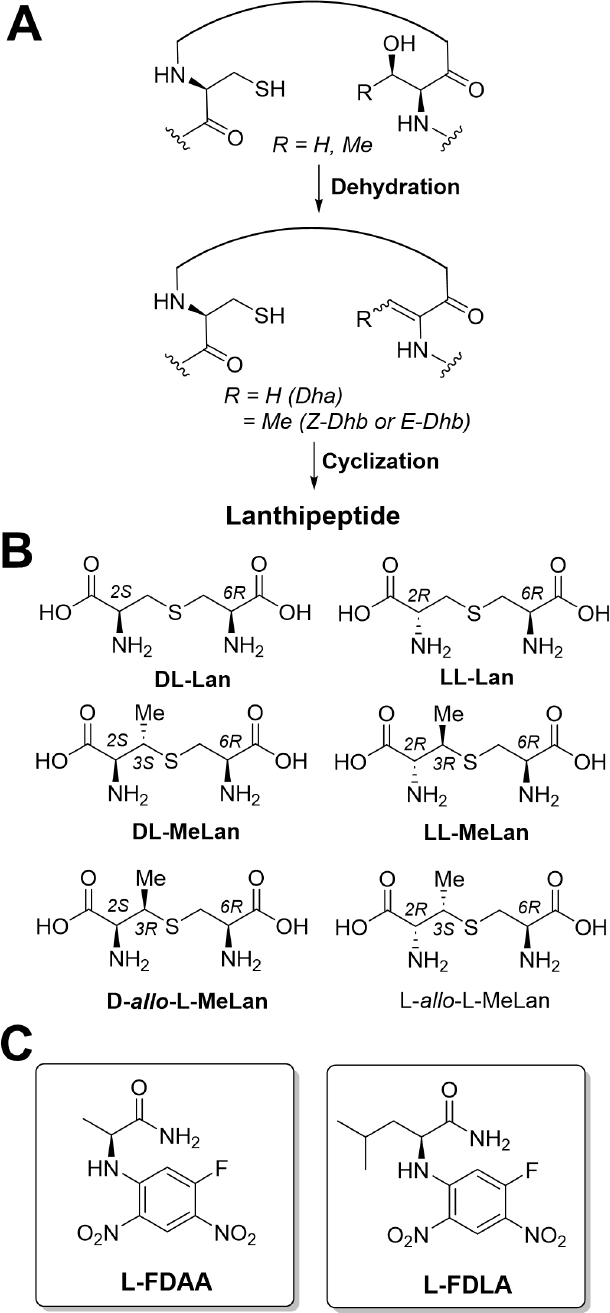
(A) Lanthipeptide synthases catalyze the formation of lanthionines and methyllanthionines by dehydrating serine/threonine residues to generate dehydroalanine (Dha) and dehydrobutyrine (Dhb), followed by the thiol-Michael addition of L-cysteine onto the Dha/Dhb moieties. (B) Theoretical diastereomers of two lanthionines (Lan) and four methyllanthionines (MeLan). Five bolded diastereomers have been discovered in lanthipeptides biosynthesis, L-*allo*-L-MeLan has thus far not been reported in natural lanthipeptides. (C) Structures of L-FDAA and L-FDLA.

Presently, researchers have identified five (Me)Lan diastereomers in characterized lanthipeptides, comprising two configurations of Lan, (2*S*,6*R*) and (2*R*,6*R*), and three stereoisomers of MeLan, (2*S*,3*S*,6*R*), (2*R*,3*R*,6*R*), and (2*S*,3*S*,6*R*).^6-10^ These isomers have been named DL- and LL-Lan, and DL-, LL-, and D-*allo*-L-MeLan, respectively (**Figure 1**). Their configurations were determined by first hydrolyzing the peptides to their individual amino acids and then chemical manipulation to obtain volatile derivatives. These compounds were then analyzed by gas chromatography (GC) monitored by mass spectrometry using a column with a chiral stationary phase (CP-chirasil-L-Val) and compared to synthetic standards obtained by a multi-step process.^8,11^ These methods have been successful but require relatively large amounts of material (∼1-2 mg for a typical lanthipeptide), a gas chromatograph, a chiral column, and chemical synthesis of standards of defined stereochemistry. Furthermore, in our hands the chemical derivatization of the amino acids and subsequent injection onto the GC column damages the column that has a considerable price-tag (∼$1400) such that its use is severely shortened (typically about 1 year of use). An alternative method that has been used is chemical desulfurization of Lan/MeLan residues and determination of the stereochemistry of the resulting Ala and 2-aminobutyric acid,^12^ but this strategy will not report on the stereochemistry on carbon 3 of MeLan.

Studies have demonstrated that altering the stereochemistry of the (Me)Lan residues in the lanthipeptide lacticin 481 leads to the abolishment of its bioactivity.^13^ This observation underscores the critical role of stereochemistry in the biological function of lanthipeptides. Whereas it was once assumed that lanthipeptides all contained Lan/MeLan with the DL stereochemistry, recent studies have demonstrated that the stereochemistry of these residues in natural lanthipeptides is much more diverse and not always readily predicted.^9,10,14-18^ Investigating the structural diversity and the interplay of stereochemistry in lanthipeptides holds significant importance in understanding their mechanism of action and exploring their application. But the stereochemistry of the Lan and MeLan residues of the great majority of newly reported lanthipeptides is not determined. Therefore, as more and more lanthipeptides are discovered by genome mining,^19-45^ a convenient method for determining their stereochemistry that is accessible to most laboratories would be valuable. We report here such a method including access to standards that use biochemical approaches that are complementary to chemical synthesis and that use methods available in most if not all laboratories studying lanthipeptides or RiPPs.

Marfey’s reagent, 1-fluoro-2,4-dinitrophenyl-5-L-alanine amide (L-FDAA, **Figure 1C**), has been widely used in peptide structure determination due to its ability to provide a robust and reliable method for analyzing the stereochemistry of amino acids in complex mixtures.^46,47^ The process involves hydrolysis of the peptide sample under acidic conditions, followed by derivatization of the amino groups with 1-fluoro-2-4-dinitrophenyl-5-L-alanine amide (FDAA, **Figure 1C**) to form diastereomers. Subsequently, these diastereomers are separated by high-performance liquid chromatography (HPLC). By analyzing the elution order and retention time and comparing them with synthetic standards, the absolute stereochemistry of the hydrolyzed fragments in the peptide sample can be determined.

The increase in the number of possible derivatizations originating from multiple reactive groups such as amino, hydroxyl, and thiol groups in an amino acid analog can pose a challenge to Marfey’s analysis.^47^ Given that Lan and MeLan are bis amino acids, they contain two sites of derivatization (**Figure 1**). The resulting problem was exemplified in the determination of the stereochemistry of D-*allo*-L methyllanthionine.^10^ In this case, bis-derivatization of the two amino groups with L-FDAA resulted in overlap of the peaks of derivatized DL-MeLan and D-*allo*-L MeLan in extracted ion chromatography (EIC).^10^ One possible explanation for this overlap is the limited interaction of FDAA with diastereomers that differ only in the stereochemistry at position 3 (**Figure 1**). We hypothesized that increasing the steric effect on the derivatization reagent might lead to full separation of the peaks of such diastereomers. In this regard, the use of 1-fluoro-2,4-dinitrophenyl-5-L-leucine amide (L-FDLA, **Figure 1C**) has been considered as an alternative approach, as it has been reported a more robust reagent for amino acids characterization.^48^ Use of FDLA is often referred to as Advanced Marfey’s analysis. We show herein that FDLA can distinguish the five known stereoisomers of (Me)Lan, and we describe how standards of these isomers can be obtained using publicly available resources. We illustrate the use of the methods with lanthipeptides discovered by genome mining.

## EXPERIMENTAL SECTION

### Lan and MeLan standard preparation

#### Nisin

Nisin was obtained by extraction from a Sigma-Aldrich product that contains about 5% of nisin from *Lactococcus lactis*. The process involved suspending 100 mg of the commercial product in 10 mL of extraction solvent consisting of 80% MeCN/20% H_2_O + 0.1% trifluoro acetic acid (TFA) in a 50 mL conical tube, followed by vigorous vortexing. After centrifuging at 4,000 xg for 10 min, the supernatant was collected, and the extraction of the insoluble material was repeated twice with 10 mL of the same solvent. The combined supernatant was then filtered using a 10K Amicon® Ultra-15 Centrifugal Filter, and the resulting flow-through was collected. The solution was frozen with liquid nitrogen and lyophilized. The resulting dry solid was redissolved in 10% MeCN / 90% H_2_O + 0.1% TFA, filtered, and injected onto an Agilent preparative HPLC system for purification using a Phenomenex Luna C5 column (10 μm, 250 × 10 mm). The following gradient conditions were used for the purification process: flow rate 4 mL/min; solvent A H_2_O + 0.1% TFA, solvent B MeCN + 0.1% TFA; 0-5 min with 2% B, 5-15 min using a gradient of 2-40% B, 15-30 min using a gradient of 40-100% B, with nisin eluting at approximately 20-22 min, corresponding to around 40% solvent B.

#### mCylL_L_ and mCylL_S_

The modified cytolysin L precursor peptide was expressed in *E. coli* BL21 (DE3) cells using the plasmid pRSFDuet-1/CylL_L_/CylM-2 (Addgene ID #208759), following the method previously reported.^9^ For the preparation of lanthionine standard, 1 L of expression of mCylLs in LB medium was typically sufficient.

Purification was modified to ensure sample purity and prevent protease digestion. After immobilized metal affinity chromatography (IMAC) purification, the elution was subjected to further purification using an Agilent preparative HPLC with a Phenomenex Luna C5 column (5 μm, 250x10 mm). The following gradient conditions were used for the purification process: flow rate 4 mL/min, solvent A H_2_O + 0.1% TFA, solvent B MeCN + 0.1% TFA; 0-5 min with 2% B, 5-15 min using a gradient of 2-50% B, 15-30 min using a gradient of 50-60% B, 30-35 min using a gradient of 60-100% B. mCylL_L_ eluted at 20-24 min, corresponding to 46-47% B.

The collected fractions were lyophilized to dryness, and then redissolved in 10% MeCN / 90% H_2_O + 0.1% TFA before injection onto an Agilent analytical HPLC system for purification using a Thermo Scientific C8 Hypersil GOLD column (5 μm, 4.6 × 250 mm). The following gradient conditions were used for the purification process: flow rate 1.0 mL/min, solvent A H_2_O + 0.1% TFA, solvent B MeCN + 0.1% TFA. The gradient profile was as follows: 0-2 min with 2% B, 2-10 min using a gradient of 2-30% B, 10-15 min using a gradient of 30-40% B, and 15-20 min using a gradient of 40-100% B. mCylL_L_ eluted at 11-12 min, corresponding to 32-35% B. To ensure purity, only fractions showing significant absorption at wavelengths of 254 nm and 280 nm on the diode-array detection (DAD) were collected and lyophilized for further usage. This method is also applicable to the expression and purification of mCylL_S_ (plasmid available at Addgene ID #208760).

#### mCoiA_1_

The modified mCoiA_1_ was expressed in *E. coli* BL21 (DE3) cells using the plasmids pRSFDuet-1 His_6_-SUMO-CoiA1_CoiB and pETDuet-1 Coi_CoiSA_(ED)_ (Addgene ID #208761 and #208762), following the method previously reported.^10^ For the preparation of methyllanthionine standard, 1 L of expression of mCoiA_1_ in TB medium was typically sufficient.

To ensure the purity of mCoiA_1_, the lyophilized elution from preparative HPLC purification was redissolved in 10% MeCN / 90% H_2_O + 0.1% TFA, and subjected to Agilent analytical HPLC purification using a Vydac C18 column (5 μm, 250 mm, No. 218TP54). The following gradient conditions were used for the purification process: flow rate 1.0 mL/min, solvent A H_2_O + 0.1% TFA, solvent B MeCN + 0.1% TFA. The gradient profile was as follows: 0-2 min with 2% B, 2-10 min using a gradient of 2-60% B, 10-15 min using a gradient of 60-80% B, and 15-20 min using a gradient of 80-100% B. mCoiA_1_ eluted at 13-15 min, corresponding to 70-80% B. To ensure purity, only fractions showing significant absorption at 254 nm were collected and lyophilized for further usage.

#### mBuvA

Plasmids were constructed and used to transform *E. coli* using previous reported method.^25^ Single colonies were selected and amplified in 5 mL of starter cultures in LB containing 50 μg/mL of kanamycin overnight. The starter culture was introduced into 1 L of LB medium containing 50 μg/mL of kanamycin and incubated at 37 °C with continuous shaking at 220 rpm until it reached an OD600 of approximately 0.8. Subsequently, the cultures were cooled on ice for approximately 15 min before the addition of isopropylthio-β-galactoside (IPTG) to a final concentration of 0.5 mM. Following this, the cultures were shaken overnight at 180 rpm at 18 °C. Cells were harvested by centrifugation at 6000 x *g* for 15 min at 4 °C. The supernatant was discarded, and the pellet was stored at -80 °C before purification.

The purification method was identical to that described for mCylL_L_, followed by an Agilent preparative HPLC step using a Phenomenex Luna C5 column (5 μm, 250x10 mm). The following gradient conditions were used for the purification process: flow rate 4 mL/min, solvent A H_2_O + 0.1% TFA, solvent B MeCN + 0.1% TFA; 0-5 min with 2% B, 5-35 min using a gradient of 2-80% B, 35-40 min using a gradient of 80-100% B. mBuvA eluted at 27-32 min, corresponding to 62-72% B.

### Peptide hydrolysis and derivatization of amino acids

The peptide sample (0.02-0.1 mg) was added to a PYREX® 15 mL screw cap glass tube, followed by 0.8 mL of 6 M DCl in D2O. N2 was bubbled through the solution for 1 min, then the tube was immediately sealed, and the mixture was stirred at reflux at 120 °C for 20 h or 155 °C for 3 h (see SI Fig.1). The resulting mixture was dried using a rotary evaporator with the water bath over 80 °C, and subsequently re-dissolved in deionized water and dried twice more. Next, 0.6 mL of 0.8 M NaHCO_3_ (in H_2_O) and 0.4 mL of 10 mg/mL L-FDLA or D-FDLA (in MeCN) were added. Following stirring in the dark at 67 °C for 3 h, 0.1 mL of 6 M HCl was gently added and the mixture vortexed vigorously. The mixture was then frozen using liquid nitrogen and lyophilized in the dark until dry. The dried solid was redissolved in 1 mL of MeCN and vortexed vigorously. The suspension was carefully transferred into 1.5 mL Eppendorf tubes using a long glass pipette and centrifuged at 16,000 x*g* for 15 min. Afterward, the resulting supernatant was transferred to screw vials for LC-MS analysis. For samples with low peak intensity of (methyl)lanthionines, the extraction solution was further concentrated up to 5-fold before reanalysis.

### LC-MS analysis

LC-MS analysis was performed on an Agilent 6545 LC/Q-TOF instrument. A Kinetex F5 Core-Shell HPLC column (1.7 μm F5 100 Å, LC Column 100 × 2.1 mm) was used to analyze reactions of Lan/MeLan with L-FDLA and D-FDLA, respectively. During the analysis, the temperature of all columns was maintained at 45 °C. HPLC parameters were: flow rate 0.40 mL/min, and use of two mobile phases (A: H_2_O+0.1% formic acid; B: acetonitrile). The gradient was set as: 2%-30% B over 0-2.5 min, 30%-80% B over 2.5-10 min, 80%-100% B over 10-10.5 min, 100%-2% B over 10.5-11 min, then 2% B re-equilibrium for 2.5 min. The mass spectrometer was set to ion polarity (negative mode), dual AJS ESI (gas temperature 325 °C, drying gas 10 I/min, nebulizer 35 psi, sheath gas temperature 375 °C, sheath gas flow rate 11 L/min, VCap 3500 V, nozzle voltage 0 V), MS TOF (Fragmenter 125 V, skimmer 65 V, Oct 1 TF Vpp 750 V) and acquisition parameters (automatic MS/MS mode, mass range 50-1700 m/z). To reduce potential contamination of the ion source by high concentrations of unreacted derivatization reagent (FDLA, elution time ∼6.5 min), a volume of 2-5 μL per injection is recommended. To avoid potential contamination of the ion source by large amounts of unreacted L-FDAA, the liquid chromatography stream was injected into the mass spectrometry starting at 7.0 min as the L-FDAA eluted around 6.7 min. Any remaining LC eluent was then directed into waste, without further MS analysis.

## RESULTS AND DISCUSSION

### Advanced Marfey’s analysis of Lan

To validate the effectiveness of FDLA and optimize the method, we initially selected two representative and easily accessible lanthipeptides, nisin and cytolysin L. Nisin is a commercially available antibacterial peptide derived from *Lactococcus lactis* and is commonly used as a food preservative (**Figure 2A**).^49-51^ It contains one DL-Lan and three DL-MeLan.^6,11^ We obtained a commercial nisin extract from *Lactococcus lactis* and purified it using HPLC. After acidic hydrolysis, the results of FDLA derivatization showed the expected product peaks by EIC that were observed in significantly higher intensity (approximately 5-10-fold) compared to FDAA treatment of the same sample (**Figure 2A**).

**Fig. 2.**
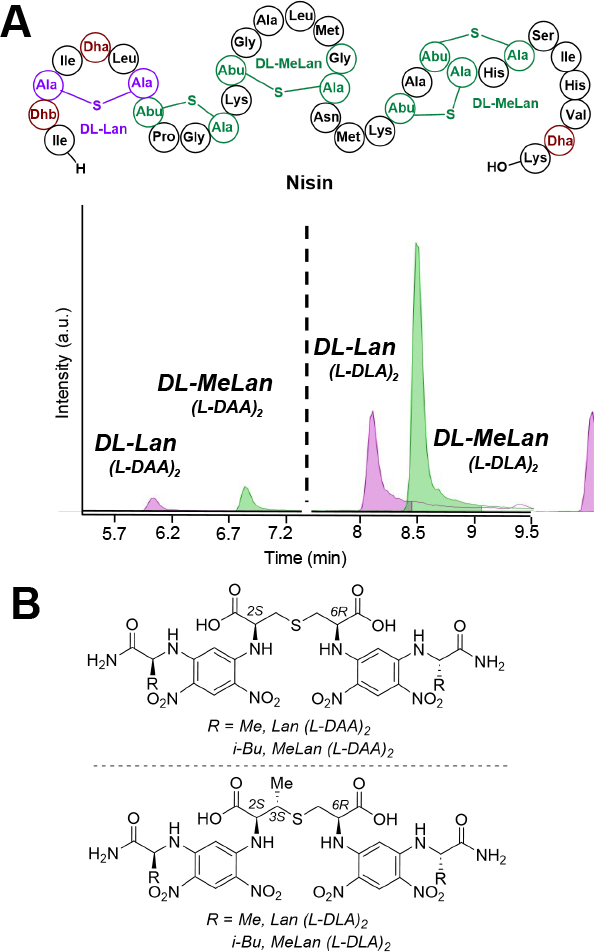
(A) Nisin structure and comparison of extracted ion chromatograms after derivatization of hydrolyzed nisin with either L-FDLA or L-FDAA. The masses monitored were DL-Lan (L-DAA)_2_ at [M-H]^-^ m/z = 711.1434 Da, DL-MeLan (L-DAA)_2_ at [M-H]^-^ m/z = 725.1591 Da, DL-Lan (L-DLA)2 at [M-H]^-^ m/z = 795.2373 Da, and DL-MeLan-(L-DLA)_2_ at [M-H]^-^ m/z = 809.2530 Da. (B) Structures of Lan-(L-DAA)_2_, MeLan-(L-DAA)_2_, Lan-(L-DLA)_2_, and MeLan-(L-DLA)_2_.

Next, we chose the lanthipeptide cytolysin L (CylLL”),^52^ a compound that can be produced in *Escherichia coli*,^9^ and for which we deposited the plasmid at Addgene (ID #208759). Modified cytolysin L precursor peptide (mCylL_L_) comprises a three (Me)Lan system including one DL-Lan, one LL-Lan, and one LL-MeLan (**Figure 3A**).^9^ Production of mCylL_L_ in *E. coli* as a His-tagged peptide, purification using nickel affinity chromatography, and acid hydrolysis provided a mixture of amino acids. L-FDAA derivatization did not lead to full separation of the two peaks of DL- and LL-Lan while L-FDLA successfully produced two well-separated peaks of DL- and LL-Lan-FDLA adducts with evenly distributed peak area (**Figure 3B**). This observation indicates the advantages of L-FDLA over standard Marfey’s analysis, and gives the possibility for quantitative analysis of (methyl)lanthionine. Using nisin’s single DL-Lan as a standard, we elucidated the order of elution of the two diastereomers of derivatized DL- and LL-Lan (**Figure 3B**).

**Fig. 3.**
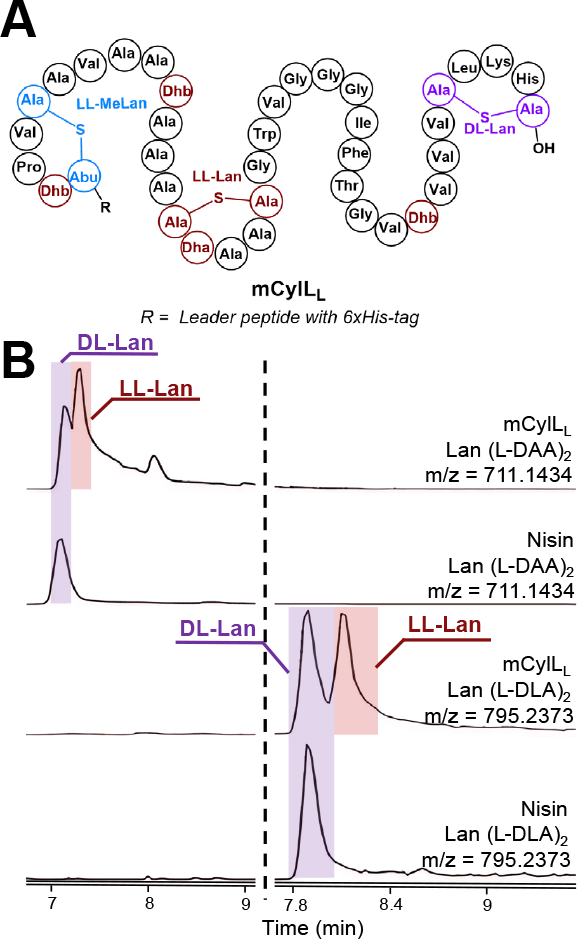
L-FDAA and L-FDLA derivatization of hydrolyzed mCylL_L_. (A) Structure of CylLL. (B) LC-MS analysis of FDAA and FDLA derivatized mCylL_L_ using a Kinetex F5 Core-Shell HPLC column. EIC monitoring of the following species: DL-Lan (L-DAA)_2_ at [M-H]^-^ m/z = 711.1434 Da, DL-MeLan (L-DAA)_2_ at [M-H]^-^ m/z = 725.1591 Da, DL-Lan (L-DLA)_2_ at [M-H]^-^ m/z = 795.2373 Da, and DL-MeLan (L-DLA)_2_ at [M-H]^-^ m/z = 809.2530 Da. Leader peptide with 6xHis-tag sequence: MGSSHHHHHHSQDPNSENLSVVPSFEELSVEEMEAIQGSGDVQAE.

Unlike most amino acids, Lan and MeLan have multiple chiral centers and therefore have a higher chance of kinetic resolution during derivatization. Derivatization with L-FDLA did not result in such resolution. The choice of MeCN/H_2_O and NaHCO3 as base in the derivatization solution proved critical to obtaining optimal intensity of the derivatized Lan/MeLan. Other derivatization conditions commonly used (e.g., acetone-triethylamine) resulted in lower peak intensities. To reduce the occurrence of the [M+Na]^+^ form, which would require integration over multiple EICs, negative mode ionization was chosen, which also resulted in intensity of the relevant ions that was consistently about 5-10 times higher compared to the analysis of [M+H]^+^/[M+Na]^+^ in the same sample.

### Advanced Marfey’s analysis of MeLan

Next, we analyzed the three previously established natural diastereomers of MeLan: DL-, LL-, and D-*allo*-L MeLan. As the MeLan standard, we selected mCoiA_1_, a modified precursor peptide that contains one of each of the three MeLan diastereomers (**Figure 4A**).^18^ Once again, the standard was prepared by expression of the His-tagged peptide in *E. coli* using plasmids deposited at Addgene (ID #208761 and #208762). The mCoiA_1_ was purified by nickel affinity chromatography, and the peptide was hydrolyzed in acid. After derivatization with L-FDLA, the LC mass spectrometry analysis showed three separate peaks (**Figure 4B**). To assign the stereochemistry of each peak, we introduced two site-directed mutations, mCoiA_1_-T43S (to remove LL-MeLan) and mCoiA_1_-T50S (to remove DL-MeLan).^10^ After applying identical treatment and analysis, the two mutations eliminated one of the three peaks each, thus assigning the stereochemistry of the corresponding MeLan and establishing the elution order of the three derivatized diastereomers. Therefore, the full-length mCoiA_1_ peptide proved to be an effective standard for characterizing all three methyllanthionine isomers (**Figure 4**).

**Fig. 4.**
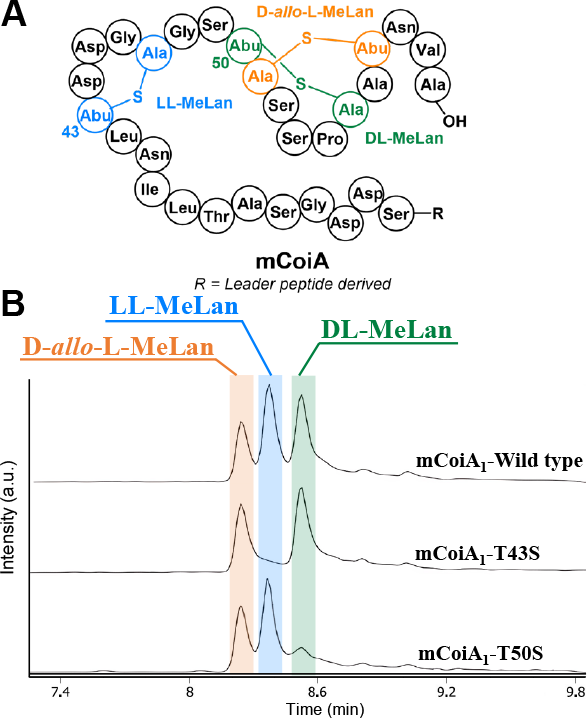
L-FDLA derivatization of hydrolyzed mCoiA_1_ and its variants. (A) Structure of mCoiA_1_. analyzed using a Kinetex F5 Core-Shell HPLC column, EIC monitoring of MeLan (L-DLA)_2_ at [M-H]^-^ m/z = 809.2523 Da. For derivatization with D-FDLA, see Figure S3. Leader peptide derived sequence: GMNANTIKGQAHSPAATAGGDAFDLDISVLE.

### Optimization and determination of the detection limit

Hydrolysis of peptides in acid at 120 °C for 20 hours is usually the standard condition for Marfey analysis, but a previous study showed that increasing the hydrolysis temperature to 150 °C for 3 hours improved efficiency without leading to breakdown or oxidation of specific residues such as Trp and Cys.^53^ Therefore, we tested these hydrolysis conditions with 0.05 mg nisin, mCylL_L_ and mCoiA_1_, and obtained similar results to those observed after hydrolysis at 120 °C for 20 h (**Figure S1**), thus significantly decreasing the time required for the analysis.

We next focused on determining the detection limit of the Advanced Marfey method for determining Lan and MeLan stereochemistry. We used the modified precursor peptide of cytolysin S (mCylLs) as model sample. This well-characterized 8 kDa, 78 amino acid sequence contains two rings, one LL-MeLan and one DL-Lan, and serves as a representative structure of a lanthipeptide genome mining exercise in that its leader peptide is still attached to the modified core peptide after expression in *E. coli*.^54^ Experimentally, we analyzed the limit of detection of mCylLs by serial dilution starting from 0.2 mg. The results suggested that 0.026 mg of the sample was sufficient to produce distinct peaks with minimal background noise (**Figure S2**). This sensitivity is a major improvement over the amount of material that is required using the GC-MS method. We also evaluated commercially available D-FDLA as derivatization agent instead of L-FDLA. The separation of the diastereomers formed upon reaction of D-FDLA with LL and DL-Lan was better than with L-FDLA, but the diastereomers formed upon reaction of D-FDLA with LL-, DL-, and *allo-*DL-MeLan provided a mixture in which two of the isomers co-eluted (**Figure S3**). Hence, L-FDLA is the reagent of choice for analysis of lanthipeptides of unknown stereochemistry.

### Application of the method to a new lanthipeptide

A previously established system known as FAST-RiPPs (<u>F</u>ast, <u>A</u>utomated, <u>S</u>calable, high-<u>T</u>hroughput pipeline for <u>RiPPs</u> discovery) combines RODEO,^55^ a tool for mining genomes to identify specific RiPP biosynthetic gene clusters (BGCs) of interest, with an automated pathway refactoring platform,^56^ and employs *E. coli* for heterologous RiPP expression.^28^ FAST-RiPPs was successful in producing a large number of novel lanthipeptides, but the stereochemistry was determined for only a subset of them because of the aforementioned need of 1-2 mg of purified final product, which could not always be obtained. In this study, we chose one such compounds for which the stereochemistry is not known, LanII-21. This compound is the product of the *buv* BGC identified in the genome of *Butyrivibrio* sp. VCD2006, butyrate-producing bacteria inhabiting the rumens of ruminant animals. The *buv* BGC encodes multiple proteins, including a rSAM BuvX (WP_026528692.1), a class II lanthipeptide synthetase BuvM (WP_081674439.1), a cyclase BuvC (WP_081674440.1), and a precursor peptide BuvA (WP_026528694.1). These genes were incorporated into a pET28a plasmid backbone for heterologous expression and post-translational modification in *E. coli* (**Figure 5A**). After purification, 0.05 mg of mBuvA was hydrolyzed at 150 °C for 3 hours, followed by derivatization with L-FDLA and LC-MS analysis. The result shows a single DL-Lan peak compared to the Lan-standard obtained from mCylL_L_, as confirmed by co-elution upon co-injection with the standard (**Figure 5B**). No significant MeLan (L-DLA)_2_ peak was obtained. This observation suggests that mBuvA forms a 5-membered DL-Lan ring at the C-terminus since that is the only location to form a Lan. To further confirm this structure, we digested mBuvA with AspN and performed tandem ESI-MS analysis, which resulted in no fragments being observed within the proposed lanthionine ring (**Figure 5C**), consistent with our assignement.

**Fig. 5.**
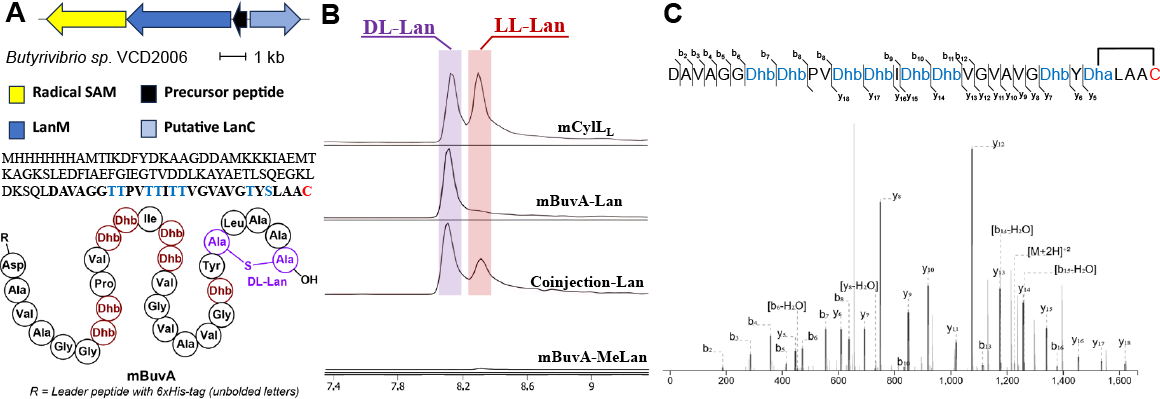
(A) The *buv* BGC. Listed are the producing organism, gene diagram, precursor peptide sequence with the predicted structure based on the result of stereochemical and LC-MS/MS analysis. Leader peptide with 6xHis-tag sequence: MHHHHHHAMTIKDFYDKAAGDDAMKKKIAEMTKAGKSLEDFIAEFGIEGTVDDLKAYAETLSQEGKLDKSQL. (B) L-FDLA derivatization of hydrolyzed mBuvA using the standard treatment procedure used herein. (C) Tandem ESI-MS of mBuvA after proteolysis with AspN (resulting in the modified peptide shown in bold font in panel A). For theoretical and observed *m/z*, see **SI Table S1**.

## CONCLUSIONS

In this study, we used L-FDLA and D-FDLA as derivatization reagents for establishing the stereochemistry of the bis-amino acids lanthionine and methyllanthionine. We used two sets of plasmids that we have deposited in a publicly accessible plasmid collection to make two samples that contain Lan and MeLan with all five previously reported configurations. Using these standards, we report the elution order of all isomers bis-derivatized with L-FDLA and D-FDLA using LC-MS. We show that as little as 0.05 mg of a lanthipeptide with its leader peptide still attached is sufficient for determining the stereochemistry of its Lan and MeLan, which is a considerable improvement of the GC-MS method that we and others have used. This approach is advantageous for cases where limited material is available and for laboratories that have molecular biology expertise but not synthetic chemistry capabilities. As such, this work provides a method that is complementary to GC-MS with synthetic standards.

## ASSOCIATED CONTENT

### Supporting Information

The Supporting Information is available free of charge at: Supporting figures

### Author Contributions

Y.L. and S. X. conceptualization, experimental data acquisition and manuscript writing. W.A.V. conceptualization and manuscript writing. All authors have given approval to the final version of the manuscript.

### Notes

The authors declare no competing financial interest.

## ACKNOWLEDGMENTS

This study was supported by a grant from the National Institutes of Health (Grant R01 AI144967 to W.A.v.d.D.) and the Howard Hughes Medical Institute. This study is subject to the Open Access to Publications policy of the Howard Hughes Medical Institute (HHMI). HHMI laboratory heads have previously granted a nonexclusive CC BY 4.0 license to the public and a sublicensable license to HHMI in their research articles. Pursuant to those licenses, the author-accepted manuscript of this article can be made freely available under a CC BY 4.0 license immediately upon publication.

