## Supporting Figures for "A Facile Method for Determining Lanthipeptide Stereochemistry"

Table of Contents

Hydrolysis temperature evaluation ………………………………………………………….. S2

Limitation of Detection (LOD) analysis ..……………………………………………………… S3

D-FDLA derivatization …...……………………………………………………………………….. S4

Fast-RiPPs LC-MS/MS Analysis ……………………………………………………..…… S5

**Hydrolysis temperature evaluation**

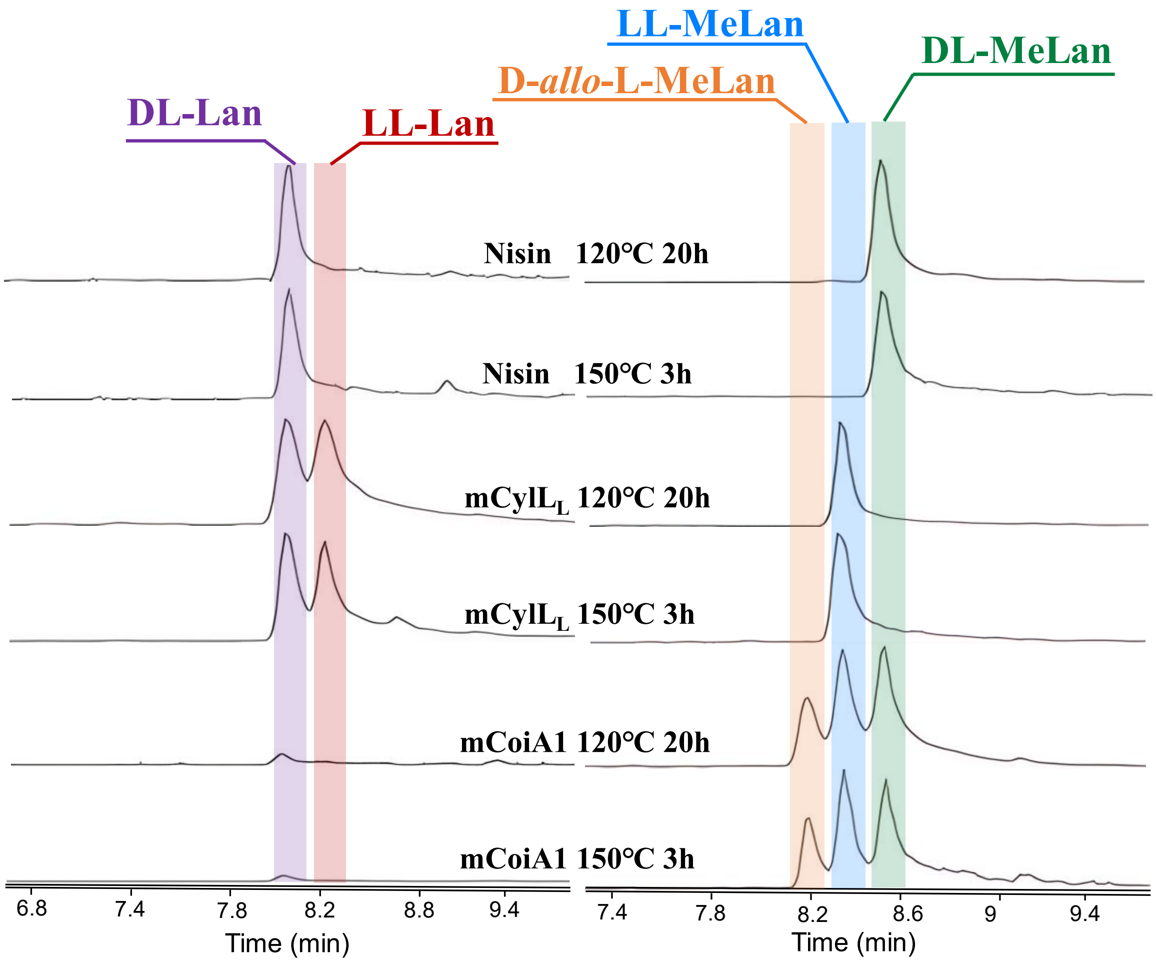


Figure S1. Influence of hydrolysis temperature on L-FDLA derivatized Nisin, mCylL$_L$ and mCoiA1. Shorter hydrolysis times at higher temperatures do not result in a change in outcome but greatly reduce the time required to determine stereochemistry as hydrolysis, derivatization and LC analysis can all be completed in one day.

### Limit of Detection (LOD) analysis

mCylLs samples were subjected to a 1.5-fold dilution in a solvent mixture of 50% MeCN and 50% $H_2O$, then transferred into two sets of PYREX® 15 mL screw cap glass tubes and dried by lyophilization. Subsequently, the 12 samples underwent standard hydrolysis and derivatization procedures as described in the main text. One set of samples was derivatized using L-FDAA, while the other derivatized with L-FDLA. Next, all samples were analyzed by LC-MS with a Kinetex F5 Core-Shell HPLC column (1.7 µm F5 100 Å, LC Column 100 x 2.1 mm). The results are shown below.

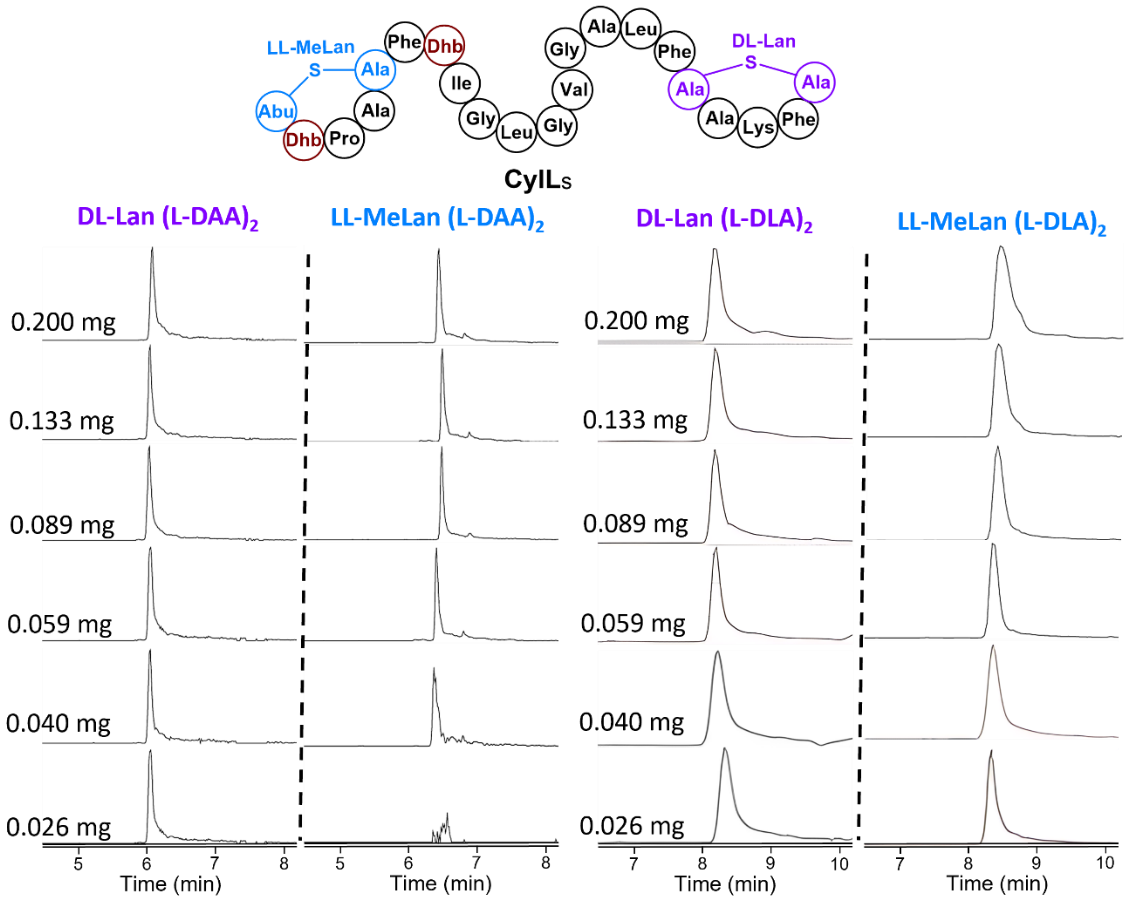


Figure S2. Limit of Detection analysis of L-FDAA and L-FDLA derivatization of hydrolyzed mCylLs using LC-MS as described in the main text.

**D-FDLA derivatization test**

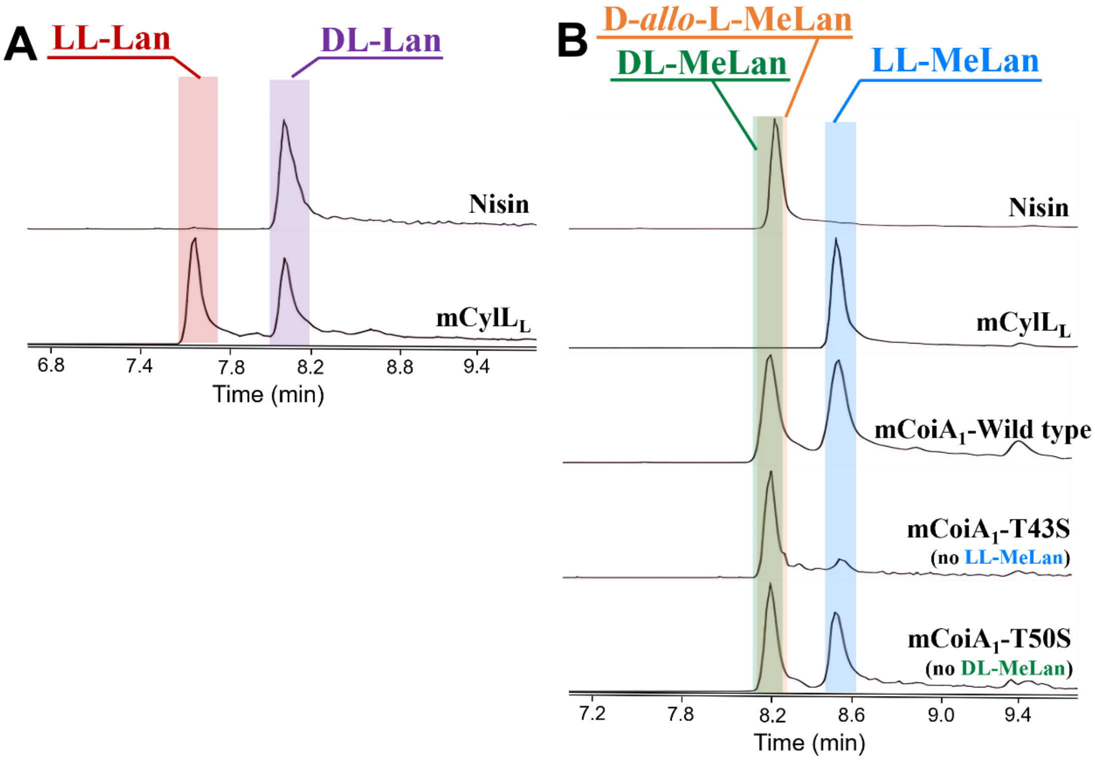


Figure S3. D-FDLA derivatized hydrolyzed nisin, mCylL$_L$, mCoiA1 and its variants. (A) EIC monitoring of Lan (L-DLA)$_2$ at $[M\text{-}H]^-$ m/z = 795.2373 Da. (B) EIC monitoring of MeLan (L-DLA)$_2$ at $[M\text{-}H]^-$ m/z = 809.2530 Da. The derivatized D-*allo*-L-MeLan and DL-MeLan isomers co-elute suggesting that L-FDLA is the preferred derivatization agent.

### Fast-RiPPs LC-MS/MS Analysis

mBuvA (100 µM, 50 uL) was digested with 5 µM AspN in 50 mM Tris-HCl, 2.5 mM $ZnSO_4$, pH in 37 °C for 4 hours. After the incubation, the sample was injected into LC-MS/MS with a AdvanceBio Peptide Plus (2.7 µm particle size, 150 x 2.1 mm) column.

SI Table S1. Theoretical and observed *m/z* ratios of AspN-digested mBuvA fragments (ions annotated in Fig. 5C)

| Ion | Theoretical *m/z* | Observed *m/z* | Mass Error (ppm) |
|---|---|---|---|
| $b_2$ | 187.0713 | 187.0710 | -1.9804 |
| $b_3$ | 284.1397 | 284.1397 | -0.3301 |
| $b_4$ | 357.1769 | 357.1770 | +0.3683 |
| $b_5$ | 414.1983 | 414.1977 | -1.4304 |
| $b_6$ | 471.2198 | 471.2199 | +0.3470 |
| $b_7$-$H_2O$ | 536.2463 | 536.2442 | -3.8921 |
| $b_7$ | 554.2569 | 554.2563 | -1.0677 |
| $b_8$ | 637.2940 | 637.2938 | -0.3093 |
| $b_{10}$ | 833.4152 | 833.4147 | -0.5941 |
| $b_{14}$-$H_2O$ | 1177.6000 | 1177.5964 | -3.0540 |
| $[M+2H]^{2-}$ | 1226.6201 | 1226.6182 | -1.5772 |
| $b_{15}$-$H_2O$ | 1260.6372 | 1260.6239 | -3.4044 |
| $y_5$ | 446.2068 | 446.2071 | +0.7990 |
| $y_6$ | 609.2701 | 609.2696 | -0.8283 |
| $y_7$ | 692.3072 | 692.3068 | -0.5905 |
| $y_8$-$H_2O$ | 731.3181 | 731.3199 | +2.4366 |
| $y_8$ | 749.3287 | 749.3283 | -0.5375 |
| $y_9$ | 848.3971 | 848.3964 | -0.7977 |
| $y_{10}$ | 919-4342 | 919.4337 | -0.5990 |
| $y_{11}$ | 1018.5026 | 1018.5017 | -0.9473 |
| $y_{12}$ | 1075.5241 | 1075.5236 | -0.4266 |
| $b_{13}$ | 1112.5735 | 1112.5766 | +2.7722 |
| $y_{13}$ | 1174.5925 | 1174.5924 | -0.0790 |
| $y_{14}$ | 1257.6296 | 1257.6289 | -0.6028 |
| $y_{15}$ | 1340.6639 | 1340.6638 | -1.1065 |
| $b_{16}$ | 1377.7161 | 1377.7154 | -0.5321 |
| $y_{16}$ | 1453.7508 | 1453.7484 | -1.6326 |
| $y_{17}$ | 1536.7879 | 1536.7853 | -1.7105 |
